# Within-patient mutation frequencies reveal fitness costs of CpG dinucleotides and drastic amino acid changes in HIV

**DOI:** 10.1101/057026

**Authors:** Kristof Theys, Alison F. Feder, Maoz Gelbart, Marion Hartl, Adi Stern, Pleuni S. Pennings

**Affiliations:** Clinical and Epidemiological Virology, Department of Microbiology and Immunology, Rega Institute for Medical Research, KU Leuven, University of Leuven, Leuven, Belgium; Department of Biology, Stanford University, Stanford, California, USA; School of Molecular Cell Biology and Biotechnology, George S. Wise Faculty of Life Sciences, Tel Aviv University, Tel Aviv, Israel; Department of Biology, San Francisco State University, San Francisco, California, USA

## Abstract

HIV has a high mutation rate, which contributes to its ability to evolve quickly. However, we know little about the fitness costs of individual HIV mutations *in vivo*, their distribution and the different factors shaping the viral fitness landscape. We calculated the mean frequency of transition mutations at 870 sites of the *pol* gene in 160 patients, allowing us to determine the cost of these mutations. As expected, we found high costs for non-synonymous and nonsense mutations as compared to synonymous mutations. In addition, we found that non-synonymous mutations that lead to drastic amino acid changes are twice as costly as those that do not and mutations that create new CpG dinucleotides are also twice as costly as those that do not. We also found that G→A and C→T mutations are more costly than A→G mutations. We anticipate that our new *in vivo* frequency-based approach will provide insights into the fitness landscape and evolvability of not only HIV, but a variety of microbes.

**Author summary:** HIV’s high mutation rate allows it to evolve quickly. However, most mutations probably reduce the virus’ ability to replicate – they are costly to the virus. Until now, the actual cost of mutations is not well understood. We used within-patient mutation frequencies to estimate the cost of 870 HIV mutations *in vivo*. As expected, we found high costs for non-synonymous and nonsense mutations. In addition, we found surprisingly high costs for mutations that lead to drastic amino acid changes, mutations that create new CpG sites (possibly because they trigger the host’s immune system), and G→A and C→T mutations. Our results demonstrate the power of analyzing mutant frequencies from *in vivo* viral populations to study costs of mutations. A better understanding of fitness costs will help to predict the evolution of HIV.

## 1 Introduction

The human immunodeficiency virus (HIV) replicates with an extremely high mutation rate and exhibits significant genetic diversity within an infected host, often referred to as a “mutant cloud” or “quasispecies” [1–7]. Although mutations are crucial for all adaptive processes, they can have fitness costs. Thus, to understand the evolution of HIV, it is important to know the fitness costs of mutations *in vivo*. Fitness costs influence the probability of evolution from standing genetic variation (often referred to as pre-existing mutations). Fitness costs also determine the effects of background selection (i.e., the effects of linked deleterious mutations on neutral or beneficial mutations) and thus affect optimal recombination rates. All of these processes affect drug resistance and immune escape in HIV [8–12]. Moreover, in addition to a better understanding of evolutionary processes in HIV and in general, a detailed knowledge of mutation costs could help us discover new functional elements in the HIV genome.

In infinitely large populations, deleterious mutations are present at a constant frequency equal to *u/s*, where u is the mutation rate from wild-type to the mutant and s is the selection coefficient that reflects the negative fitness effect, or cost, of the mutation [13, 14]. In natural populations of finite size, however, the frequency of mutations is not constant; instead it fluctuates around the expected frequency of *u/s*, because of the stochastic nature of mutation and drift [13]. Due to these stochastic fluctuations of frequencies, it is impossible to accurately infer the strength of selection acting on individual mutations (i.e., their cost) from a single observation of a single (finite size) population. This is why most approaches based on the frequencies of mutations have to aggregate mutations in groups so that a distribution of frequencies (the “site frequency spectrum”) can be analyzed and compared between groups of mutations. This approach can therefore never lead to fitness estimates of individual mutations. Alternative approaches to assess fitness effects are mostly based on (1) phylogenetic or entropy-based approaches which use between-population or between-species differences (substitutions) as opposed to within-population variation [15–21] or (2) they use *in vitro* systems to measure fitness effects (e.g., times series or competition experiments in cell culture [22–26]). These approaches have their limitations. The phylogenetic approaches estimate fitness costs over very long timescales, and it is unclear how relevant those estimates are for current viral populations. The entropy-based methods focus on fairly small subsets of common mutations and exclude the vast majority of mutations because they are rare. Regarding the approaches based on *in vitro* systems, it is unclear whether fitness costs are similar to *in vivo* fitness costs.

HIV has unique properties that allow us to study fitness effects *in vivo*: It is fast evolving [27–31] and leads to persistent infections [32–34]. This means that genetic diversity accumulates quickly and independently in every host, and samples from different patients can thus be treated as independent replicate populations [35,36]. By aggregating data on the exact same mutation from many patients, the mean frequency of the mutation will approach *u/s* and can therefore be used to estimate its fitness cost, because the fluctuations in mutation frequencies represent an ergodic process [37]. Based on this logic, we present a novel approach that uses observed mutation frequencies in many HIV-infected patients to determine the fitness effects of mutations *in vivo*. For this analysis, we assume that there are no epistatic interactions and that selection coefficients and mutation rates do not vary between patients. A variation of this approach was employed in parallel to us by Zanini *et al*. to estimate HIV fitness values from nine infected patients [31]. Reassuringly our basic results overlap with Zanini *et al*.; here we also report on novel genomic insights obtained by our method.

In the current study, we demonstrate the utility of this new approach. We focus on transition mutations (A↔G and C↔T) in 870 sites of the *pol* gene, which encodes HIV’s *protease* protein and part of the reverse transcriptase (RT) protein, in 160 patients infected with HIV-1 subtype B. Transitions are much more common in HIV than transversions [29], and thus sufficient data are available for these mutations; we focus on the *pol* gene because it is highly conserved and its products experience less direct contact with the immune system than the exposed product of the much more variable envelope (*env*) gene [32,33]. Finally, we excluded mutations at drug resistance-related sites, because the samples we use came from patients receiving several different treatments. Accordingly, we expect that the mutations that we did include in our study are deleterious.

We report that this proof-of-concept of our *in vivo* frequency-based approach allowed us to quantify known properties of mutational fitness costs (such as differences between synonymous, non-synonymous and nonsense mutations), and it also revealed novel insights into the evolutionary constraints of the HIV genome (such as the surprising cost of mutations that form a CpG site and of G→A and C→T mutations). The fitness effects are surprisingly independent of the location in the gene (although we do find a small difference between mutations in *RT* versus mutations in *protease*). Because we study a large number of mutations, it was possible to determine how characteristics of mutations affected their costs in more detail than has previously been possible. Our results demonstrate the power of analyzing mutant frequencies from *in vivo* viral populations to study the fitness effects of mutations.

## 2 Results

### 2.1 Data are consistent with model assumptions

An important assumption for the proposed method is that the mutation frequencies are drawn from independent populations (each patient harbors an independent HIV population) that are in mutation-selection-drift equilibrium. This assumption could be violated if the subtype B epidemic in the United States is not in mutation-selection-drift equilibrium and if samples were taken soon after a person was infected. In that case, several patient samples may share high frequency variants of a mutation, which violates the assumption of independence. To minimize the potential confounding effect of shared high frequency variants, we removed all site/patient combinations where the mutant frequency of the sample from the first time point for a patient was not 0%. This filtering step removed 6% of the data.

A further assumption of our approach is that within-patient populations are in mutation-selection-drift balance. We tested whether the data were consistent with this assumption. For each site, we used the mean frequency of the mutant and the mutation rate estimate from Abram *et al* [29, 38] to estimate the selection coefficient. With this point estimate of the selection coefficient, the nucleotide-specific mutation rate estimate from Abram *et al* [29, 38] and a population size of N = 5, 000, we ran individual-based simulations to create 160 population frequencies for the given mutation (following [35]). Next, we sampled from these simulated populations using the sample sizes of the real data. The resulting simulated sample frequencies were then compared with the observed sample frequencies using a Mann-Whitney test. At 91% of the sites, the simulated frequencies were not significantly different than the observed frequencies, using 5% significance level. The remaining 9% may be governed by epistasis or may be adaptive, or may have different fitness effects in different patients, so that mutation-selection balance may not describe the dynamics of these mutations well. We repeated this analysis for a range of population sizes and found very similar results (results not shown). This result gives us confidence that the mutation-selection-drift equilibrium describes the actual dynamics in the patients well (see figure 1).

**Figure 1:**
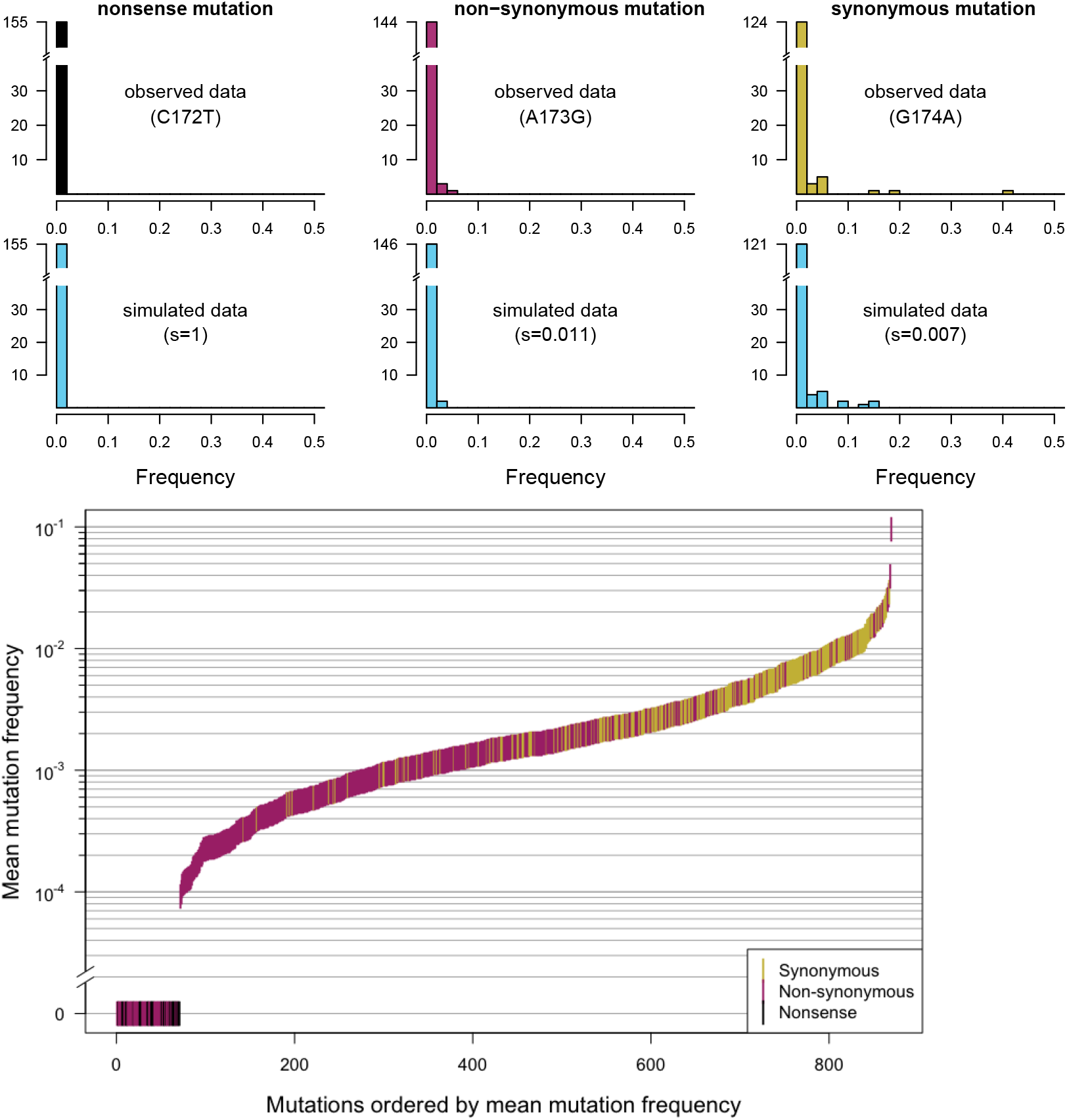
Different frequency patterns of synonymous, non-synonymous and nonsense mutations. As expected, in the HIV *pol* gene, synonymous mutations occurred more frequently than non-synonymous mutations, which occurred more frequently than nonsense mutations, which were not observed at all. A) First row: Single-site frequency spectrum for three sites in the HIV *protease* protein (sites 172, 173 and 174). Second row: simulated data based on estimated selection coefficients. B) Mean mutation frequencies for all sites, ordered by mutation frequency.

### 2.2 A clear difference between the costs of synonymous, non-synonymous and nonsense mutations

Now that we are confident that our main model assumptions hold, we compared mutation frequencies for the three main classes of mutations: synonymous, non-synonymous and nonsense mutations. As an example, we show the observed and simulated frequency spectra at all three nucleotides of codon 58 of the *protease* protein, which comprises nucleotides 172 through 174 (Fig. 1A). The transition mutation at the first position (172) creates a premature stop codon. As expected for a lethal mutation, this nonsense mutation was never observed in the data and thus has a frequency of zero in all patients. A transition mutation at the second codon position (173) leads to an amino acid change (glutamine to arginine), and also creates a CpG dinucleotide. This mutation was found at low frequencies in some patients (between 0 and 4%). The average frequency was 0.001, suggesting a selection coefficient of 0.011. A synonymous mutation at the third position of the codon (174) was observed at a wide range of frequencies (mean frequency 0.008, estimated selection coefficient 0.007, see Fig. 1A). The simulated data for all three nucleotides are shown in blue in the second row of the figure.

The pattern that synonymous mutations are found at higher frequencies than non-synonymous mutations, which were found at higher frequencies than nonsense mutations was seen in the entire dataset. To illustrate this, we ordered all sites according to observed mutation frequencies and plotted the three categories of mutations in three colors (Fig. 1B). The distributions of the mean frequencies for each of the three main categories of mutations were significantly different (one-sided two-sample Wilcoxon test, p < 2.2 · 10^-16^ for nonsense vs non-synonymous mutations and for non-synonymous vs synonymous mutations; Fig. 1B). All nonsense mutations had an average frequency of zero, and so did some non-synonymous mutations. Most non-synonymous mutations had a lower frequency than synonymous mutations (80% of non-synonymous mutations were present at a frequency lower than 0.002, whereas 82% of synonymous mutations were present at a frequency higher than 0.002). This difference in distributions probably reflects the higher cost of non-synonymous mutations, which are more likely to directly affect virus replication. This analysis therefore provides a proof of principle that our approach works: The observed frequencies reflect the relative costs we would expect for these broad categories of mutations.

### 2.3 GLM shows costs associated with mutations that create new CpG dinucleotides, G-A and C-T mutations and mutations that lead to drastic amino acid changes

To determine how various mutation characteristics affect observed frequencies of synonymous and non-synonymous mutations, we fit a generalized linear model (GLM). Nonsense mutations are excluded for this analysis because they were never observed (all frequencies were zero). The advantage of using a GLM is that we can directly analyze raw counts as opposed to frequencies. This approach automatically gives more weight to patients for whom we have more sequences, and it allows us to investigate several effects simultaneously (see Methods). The effects we considered were 1. whether a site is part of *protease* vs. reverse transcriptase, 2. the shape value (an experimentally determined measure of RNA secondary structure [39]), 3. the ancestral nucleotide (A, C, G or T), 4. whether a mutation is synonymous or non-synonymous, 5. whether a mutation would create a new CpG site and 6. whether a mutation leads to a drastic amino acid change or not. Amino acid changes were considered drastic when the transition changes the encoded amino acid from one major amino acid group (positively charged, negatively charged, uncharged, hydrophobic and special cases) to another (see Methods). The GLM results are shown in Table 1 and Fig 2. We used estimated mutation rates from Abram *et al* [29,38] and the mutation-selection formula (*f* = *u/s*) to translate the observed frequencies into selection coefficients (costs).

As we saw previously, non-synonymous mutations have lower frequencies than synonymous mutations (line 9 in Table 1, *p* < 0.001), which means that they are more costly. We will now look into synonymous and non-synonymous mutations in more detail.

**Figure 2:**
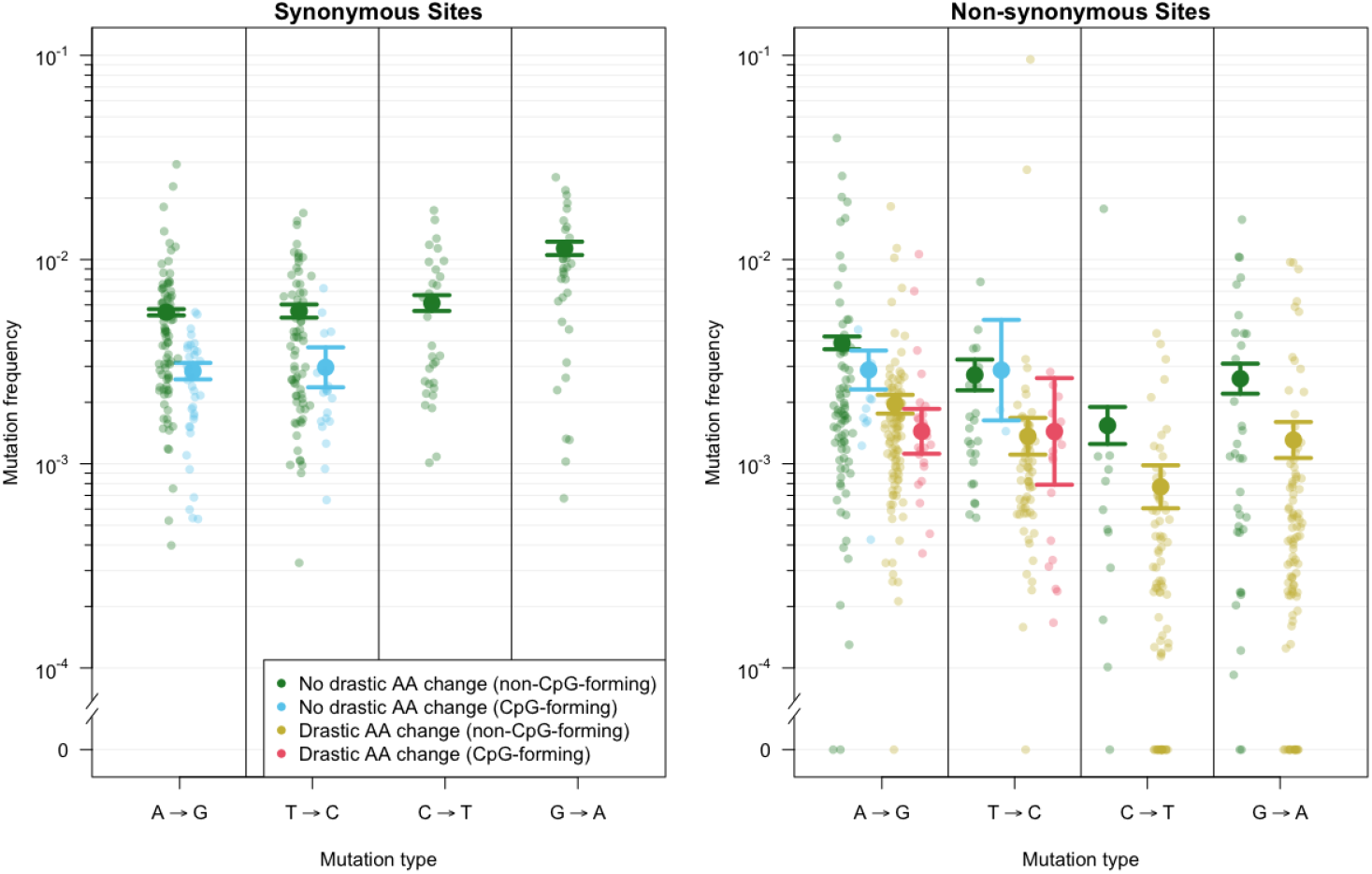
Predicted and observed mutation frequencies for different mutation classes. Mutation frequencies as predicted by the generalized linear model (large dots) and observed frequencies (small dots). The horizontal lines show the standard errors from the GLM. The graph shows the model predictions for synonymous and non-synonymous mutations that do not involve a drastic amino acid change and either form CpG sites (blue) or do not (green). In addition, for non-synonymous mutations, predictions are shown for mutations that involve a drastic amino acid change and either form CpG sites (light red) or do not (yellow).

#### CpG sites

Among synonymous mutations, a strong effect was associated with whether or not a mutation created a new CpG dinucleotide site. A→G mutations and T→C mutations that created new CpG sites were found at significantly lower frequencies than A→G mutations and T→C mutations that did not (*p* < 0.001, line 7 in Table 1) (note that G→A and C→T mutations cannot create new CpG sites). Using model-predicted frequencies and known mutation rates, we find that CpG-creating synonymous mutations are 2 times more costly (selection coefficient appr. 0.004 for both A→G mutations and T→C mutations), than the corresponding non-CpG-creating synonymous mutations (selection coefficient appr. 0.002 for both A→G mutations and T→C mutations). This finding is consistent with the hypothesis that CpG sites are costly for RNA viruses because they trigger the host antiviral cellular response [40–44].

Non-synonymous mutations that create CpG sites are also found at lower frequencies than non-synonymous mutations that do no create CpG sites (Fig. 2). However, the effect of creating a CpG site is not as strong in non-synonymous sites as it is in synonymous sites leading to a positive GLM coefficient (lines 13 and 14 in Table 1, *p* < 0.001. The difference in frequencies shows that, among mutations that do not lead to a drastic amino acid change, A→G mutations that create a CpG site are approximately one-and-a-half times more costly than those that do not (0.0039 vs 0.0028).

#### Ancestral nucleotide

We also found an effect of the nucleotide in the consensus sequence (i.e., the presumed ancestral nucleotide): synonymous G→A mutations were observed at higher frequencies than the other mutations (line 6 Table 1), but given their high mutation rate, their frequencies were actually lower than expected. We could not test whether this effect was significant using the GLM framework, but a one-sided two-sample Wilcoxon test showed that the difference in estimated selection coefficients for G→A mutations and non-CpG-forming A→G mutations was highly significant (*p* = 5 · 10^-9^). Indeed, the estimated selection coefficients based on model predictions suggested that synonymous G→A mutations are two-and-a-half times as costly as non-CpG-forming A→G mutations (0.0048 vs 0.002). For synonymous C→T mutations, their frequency is not significantly different from the frequency of synonymous, non-CpG-forming A→G mutations (see line 5 in Table 1), but because their mutation rate is about double the mutation rate of A→G mutations, their estimated cost is two times as high as for non-CpG-forming A→G mutations (0.0039 vs 0.002, *p* = 4·10^-05^), see Fig. 2 and Fig. S5. We find qualitatively similar results when we use mutation rates from [31].

**Table 1:** Predictors of frequencies for mutations in the *pol* gene, estimated using a generalized linear model (GLM). The intercept (*) is estimated for synonymous, non CpG-forming A→G mutations in *protease* with shape value 0. The predicted frequency for such mutations is therefore *e*^-5.2^ which equals 0.0055, as indicated in the last column(**). Row 2-15 of the table lists the effects of changing attributes of the mutation, which is why A→G mutations are not explicitly listed in the table. To estimate predicted frequencies for a particular class of mutations from the table, the relevant coefficient estimates must be summed, then exponentiated. For example, the predicted frequency of a synonymous, A→G mutation in *protease* with shape value 0 that would create a CpG site is *e*^-5.2-0.664^ (taken from line 7), alternatively, one could calculate this predicted frequency as 0.0055 * (1 − 0.49) = 0.0028. For a site that is CpG forming and non-synonymous, we have to add the estimates from lines 9 and 13 to get *e*^-5.2-0.664-0.345+0.358^ or 0.0055 * (1 − 0.49) * (1 − 0.29) * (1 + 0.43) = 0.0029. For the continuous shape parameter, the value of the shape parameter for a given site should be multiplied by 0.168 (line 2) and then exponentiated, e.g., for a shape value of 0.5, the predicted frequency is *e*^-5.2-0.5∗0.664^ = 0.0060.

|  |  | Estimate | Std. Error | z value | Pr (> \| z \|) | Effect |
| --- | --- | --- | --- | --- | --- | --- |
| 1 | (Intercept) | -5.199* | 0.035 | -147.037 | 0.000 | 0.0055** |
| 2 | In reverse transcriptase | 0.096 | 0.023 | 4.223 | 0.000 | +10% |
| 3 | Shape | 0.168 | 0.037 | 4.556 | 0.000 | +18% |
| 4 | T→C | 0.013 | 0.039 | 0.339 | 0.734 | +1% |
| 5 | C→T | 0.104 | 0.054 | 1.940 | 0.052 | +11% |
| 6 | G→A | 0.720 | 0.040 | 18.134 | 0.000 | +105% |
| 7 | CpG-forming | -0.664 | 0.058 | -11.520 | 0.000 | -49% |
| 8 | T→C:CpG-forming | 0.029 | 0.093 | 0.315 | 0.753 | +3% |
| 9 | Non-syn | -0.345 | 0.037 | -9.460 | 0.000 | -29% |
| 10 | T→C:Non-syn | -0.375 | 0.062 | -6.017 | 0.000 | -31% |
| 11 | C→T:Non-syn | -1.036 | 0.083 | -12.456 | 0.000 | -65% |
| 12 | G→A:Non-syn | -1.124 | 0.058 | -19.496 | 0.000 | -65% |
| 13 | Non-syn:CpG-forming | 0.358 | 0.090 | 3.995 | 0.000 | +43% |
| 14 | T→C:Non-syn:CpG-forming | 0.330 | 0.153 | 2.156 | 0.031 | +39% |
| 15 | Drastic amino acid change | -0.691 | 0.034 | -20.394 | 0.000 | -50% |

Among non-synonymous mutations, we also found a strong effect of the ancestral nucleotide: C→T and G→A mutations are both more costly than A→G and T→C mutations (Fig. 2). Again, we could not use the GLM framework to test whether this difference was significant, but one-sided two-sample Wilcoxon tests showed that the difference in estimated selection coefficients was highly significant (*p* = 4 · 10^-6^ for C→T and *p* = 3 · 10^-5^ for G→A mutations when compared with A→G mutations). We estimated that, among non-synonymous mutations that do not involve a drastic amino acid change nor create a CpG site, C→T mutations are five-and-a-half times more costly than A→G mutations (0.0157 vs 0.0028), and G→A mutations are seven times more costly than A→G mutations (0.021 vs 0.0028), see Fig. 2 and Fig. S5.

#### Drastic amino acid changes

Mutations that led to a drastic amino acid change were found at lower frequency than mutations that did not (*p* < 0.001). For example, A→G mutations that result in a drastic amino acid change are roughly twice as costly as A→G mutations that do not (0.0057 vs 0.0028). We observed similar fold changes for the other possible transitions (Fig. 2).

#### Other effects

Mutations in the *RT* portion of the gene had slightly higher frequencies than those in the *protease* portion (*p* < 0.001, line 2 in Table 1), suggesting that they are somewhat less costly. Similarly, our model predicts a small but significant effect of the shape value (*p* < 0.001, line 3 in Table 1), an experimentally determined measure of RNA secondary structure [39]. Specifically, sites with a higher shape value (i.e., those less likely to be part of an RNA structure) were associated with higher mutation frequencies (suggesting lower mutational costs), presumably because the secondary structure of the RNA molecule plays a functional role in HIV replication [39] (see Table 1).

### 2.4 Effects not captured by the GLM

#### Outliers

We asked whether we could use our results to find individual sites at which mutations are more costly than expected based on our current knowledge of the HIV genome. However, if we do a simple outlier analysis and focus on, say, the 5% most costly sites overall in our dataset, we will find that these are all the nonsense mutations, plus some mutations that lead to drastic amino acid changes and create CpG sites. Such analysis by itself is not very interesting, since our GLM analysis already revealed these results. Instead, we first grouped the sites in nine groups according to the GLM results (see Methods) and then to look at the outliers (5% highest selection coefficient values) within each of these groups. We made a table of all outliers (see Suppl. materials). We found that a few amino acids show up in the outlier list more than once, but this is not surprising, given that our dataset only comprises a few hundred amino acids. The vast majority of these sites do not have a known function; a select few are near the active site of the protein. In future work, it will be worth following up on those positions in *pol*.

#### Amino acid identity

The nature of the amino acid change (drastic or not) and the ancestral nucleotide in the consensus sequence both had an effect on costs. In addition, we found that many of the most costly non-synonymous mutations were associated with a small number of amino acid changes starting from glycine (G) and proline (P). This is consistent with our knowledge of protein structure: glycine and proline are often unique and irreplaceable, as the only cyclic and smallest amino acid, respectively. The triplets that encode these two amino acids are C and G rich (CCN for proline and GGN for glycine) which may partially explain why G→A and C→T mutations are costly. Fig 3 shows the cost of non-synonymous changes ordered by ancestral and mutant amino acid. Contrary to our results, Zanini *et al*. [31] (figure S7) found other amino acids (tryptophan (W), tyrosine (Y), cysteine (C), and also proline (P)) to contribute most to the cost, but this difference is likely due to the fact that they considered all possible mutations including nonsense mutations and synonymous mutations, which lowers the average cost of amino acids encoded by codons with synonymous mutations and increases the cost of amino acids encoded by codons with nonsense mutations. For example, tryptophan (encoded by only one codon, TGG) has no synonymous mutations and two of the three possible transitions lead to a stop codon, which makes it very costly compared to other amino acids in the Zanini analysis [31]. This might explain the discrepancy between our analyses.

#### No effect of location in the *pol* gene

We were interested to see whether fitness costs were distributed evenly along the *pol* gene or whether some parts of the gene harbored clusters of sites with particularly high cost mutations as was found by other studies [17]. We plotted the fitness cost point estimates along the length of the *pol* gene (i.e., the sites for which we have data) (see Fig. 4). We colored sites according to whether the transition mutation we considered was synonymous or non-synonymous, and the latter group was split into G-A and C-T mutations in light red and A-G and T-C mutations in dark pink. Visually, it is clear that there is no strong effect of location on fitness cost. There are no clear stretches of particularly high or low costs. We tested whether there was a statistically significant effect of location using a randomization test and we did this separately for synonymous and non-synonymous mutations using a sliding window approach (see Methods). We found no effect of location, although sites within the same codon did have correlated fitness costs.

### 2.5 Parameters for gamma distribution of fitness effects

In addition to the characteristics that determine the fitness costs of individual mutations, we investigated the distribution of fitness effects (DFE). This distribution is of interest to the evolutionary biology community because it affects standing genetic variation, background selection, and optimal recombination rates [16].

**Figure 3:**
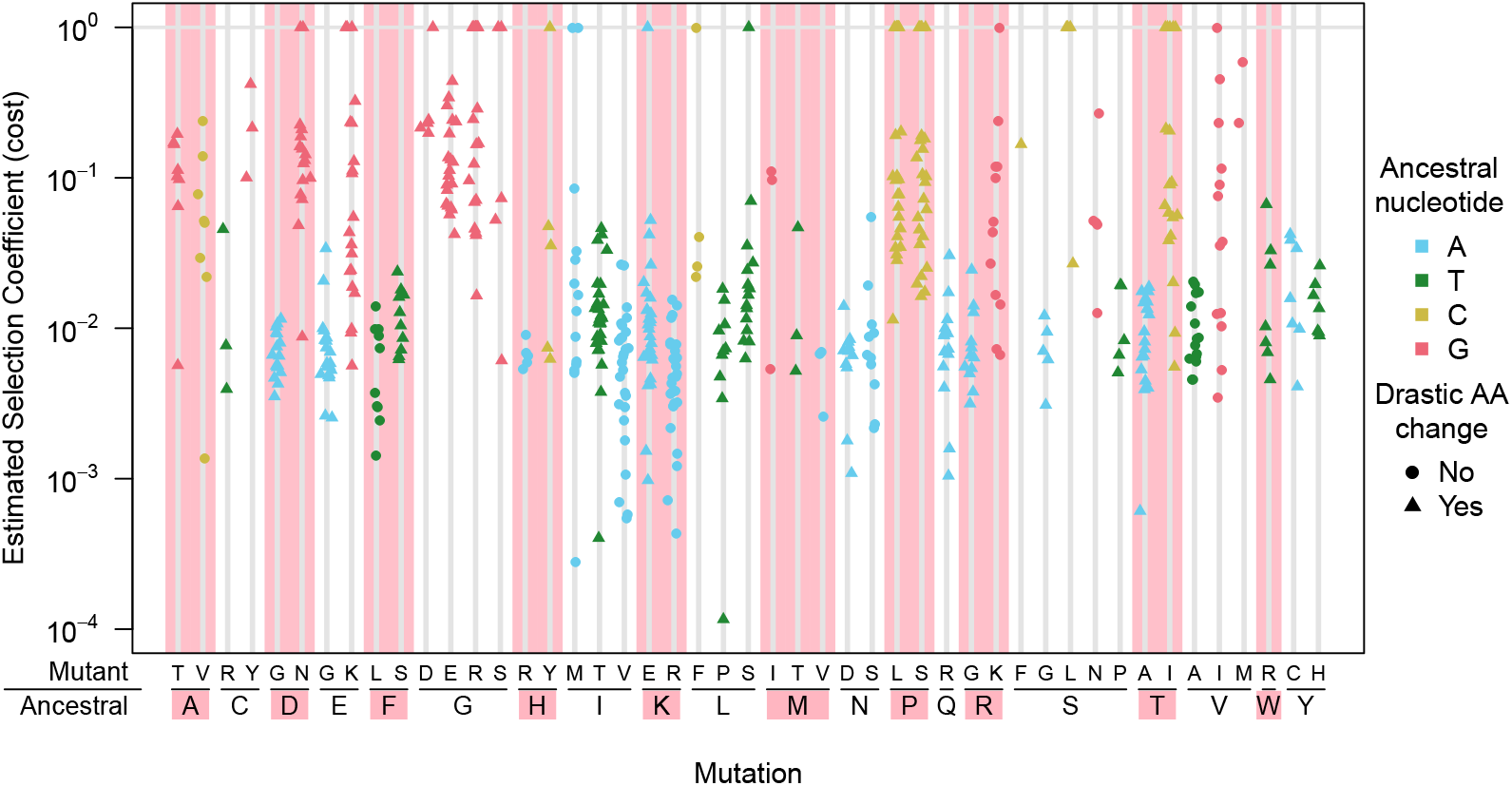
Distribution of estimated selection coefficients by amino acid replacements. Many of the most costly mutations are concentrated at a few amino acids (e.g., P (proline) and G (glycine)). The selection coefficients shown are calculated directly from mean mutation frequencies and mutation rates using the mutation-selection balance formula, *f* = *u/s*.

Moreover, the DFE affects the evolvability of a population: A DFE weighted toward neutral and adaptive mutations may reflect a population with more capacity to evolve. Many viruses, however, have been found to have a DFE composed mainly of deleterious and lethal mutations. To determine the DFE of the *pol* gene in HIV, we used the fitness cost point estimates for synonymous and non-synonymous mutations (including nonsense mutations) for each of the ancestral nucleotides (Fig. 5). Overall, there were few very deleterious and lethal mutations, except for non-synonymous C→T and G→A mutations and nonsense mutations. This is, at least partly due to the fact that we only consider transition mutations. We also estimated parameters for the gamma distribution that best describes the entire DFE (Table 2). These parameters can be used in studies of background selection and in other studies that involve simulations of evolving populations. We performed this analysis also for two other datasets with *pol* sequences for multiple patients (the Zanini dataset [45] and the Lehman [46], see Methods and suppl. materials).

-2.25in0in

**Figure 4:**
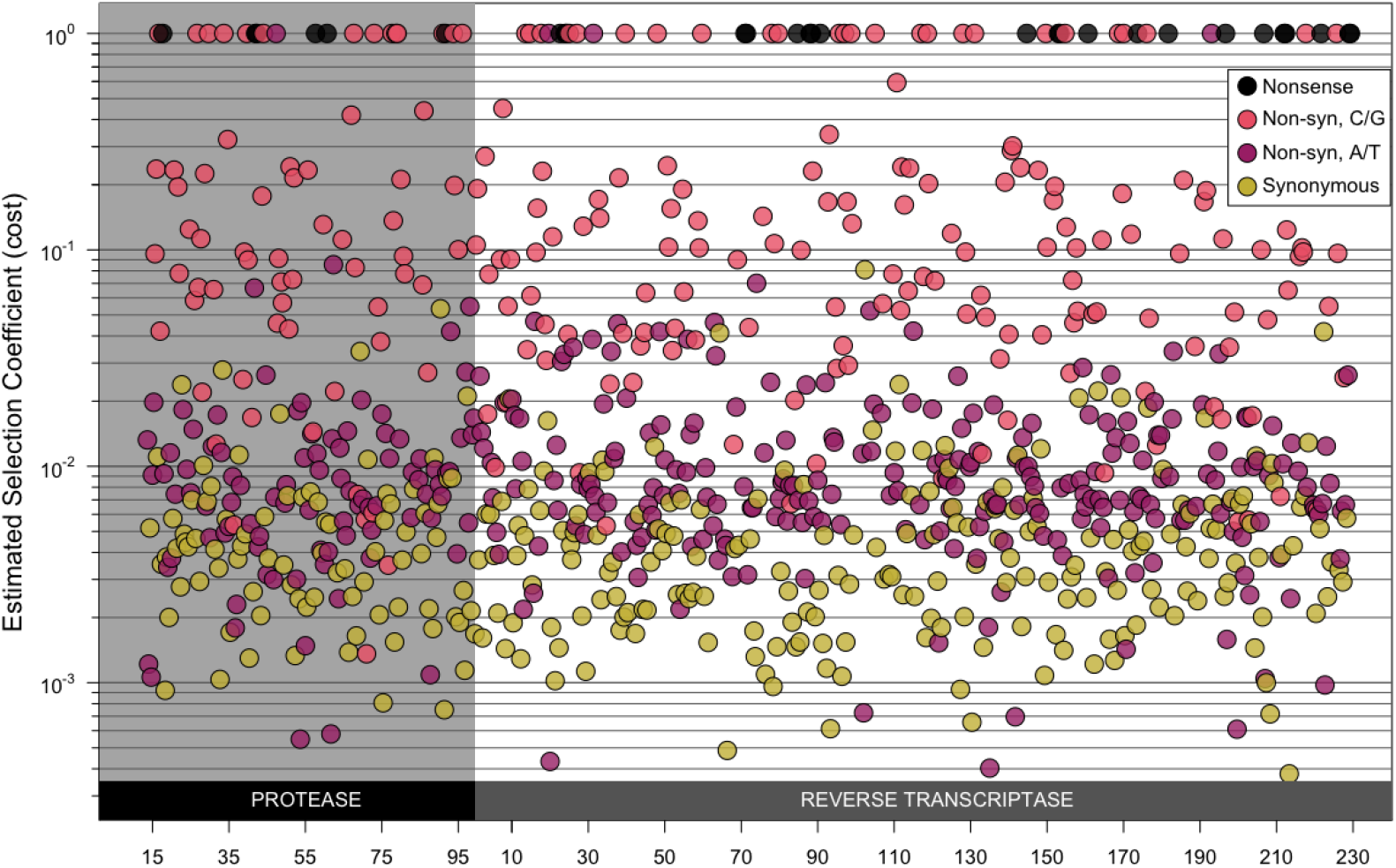
Estimated selection coefficients for transition mutations along the *pol* gene. Point estimates for the selection coefficients for each transition mutation along the *pol* gene. Synonymous mutations are shown in yellow, non-synonymous mutations are shown in light red (C→T or G→A mutations) and dark pink (T→C or A→G mutations), nonsense mutations are shown in black. This plot illustrates that estimated selection coefficients do not appear to be affected by location in the gene. Note that these histograms include mutations that create CpG sites and those that don’t, which means that the effect that G→A and C→T mutations are more costly than non-CpG forming A→G mutations is not visible in this figure.

**Figure 5:**
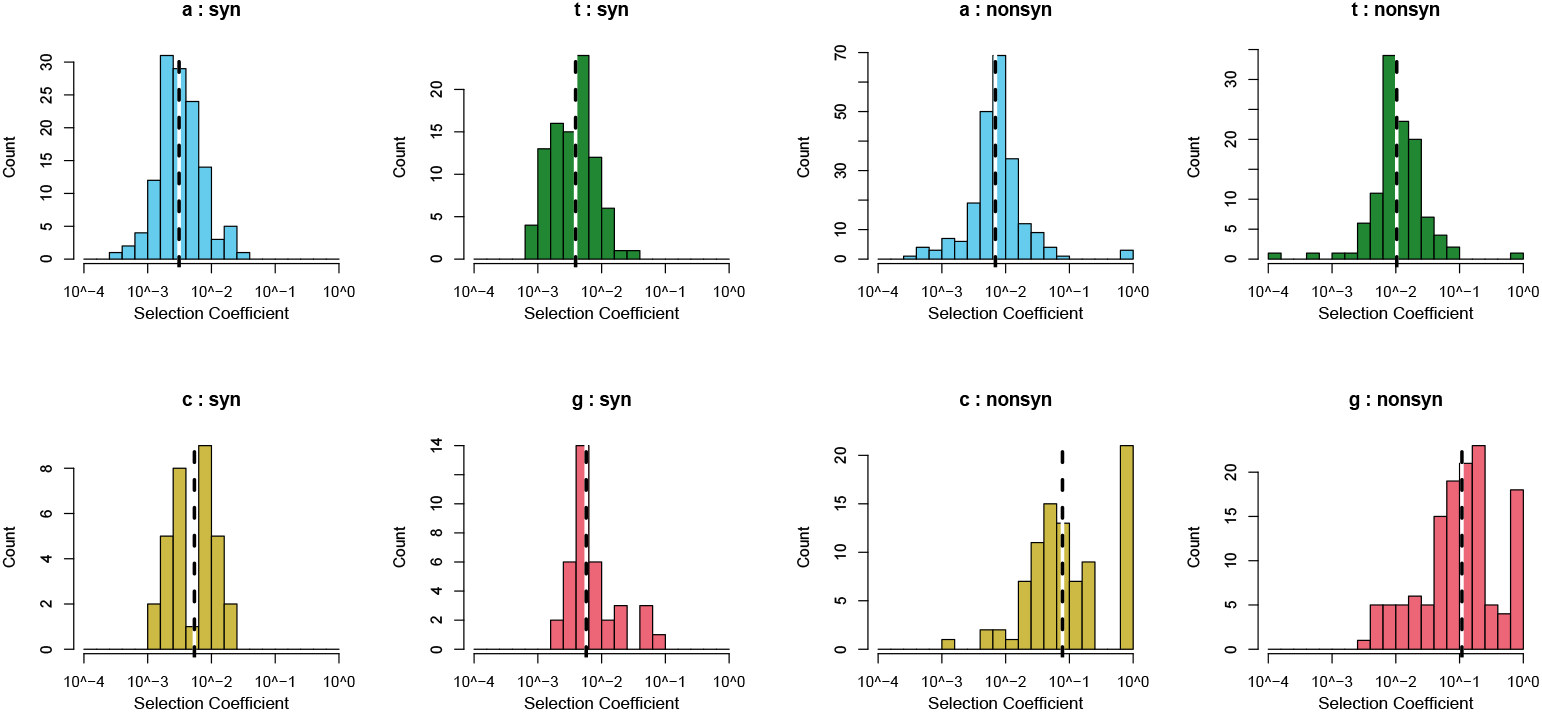
Distribution of fitness costs as estimated from mutation frequencies using the mutation-selection balance formula (*f* = *u/s*). Most synonymous mutations (left panel) have very low selection coefficients. For non-synonymous mutations (right panel), selection coefficients are higher, especially for C→T and G→A mutations. Dashed vertical lines indicate median selection coefficients. Note that the scales of the y-axes differ between the individual plots.

**Table 2:** Parameters for the gamma distribution of fitness effects for transition mutations in *pol* in 160 HIV-infected patients from the Bacheler *et al*. dataset, reflecting scale (κ) and shape *(θ)*. The ‘fraction lethal’ is the fraction of the mutations that had a mean frequency smaller than or equal to the mutation rate, so that they are estimated to be lethal.

| Num. sites | Fraction lethal | Mut Rates from Abram 2010 |  | Mut rates from Zanini 2016 |  |
| --- | --- | --- | --- | --- | --- |
| | | $\kappa$ | $\theta$ | $\kappa$ | $\theta$ |
| 870 | 0.082 | 0.334 | 0.275 | 0.327 | 0.333 |
|  | (0.066, 0.099) | (0.257, 0.411) | (0.265, 0.289) | (0.267, 0.388) | (0.321, 0.348) |

**Table 3:**
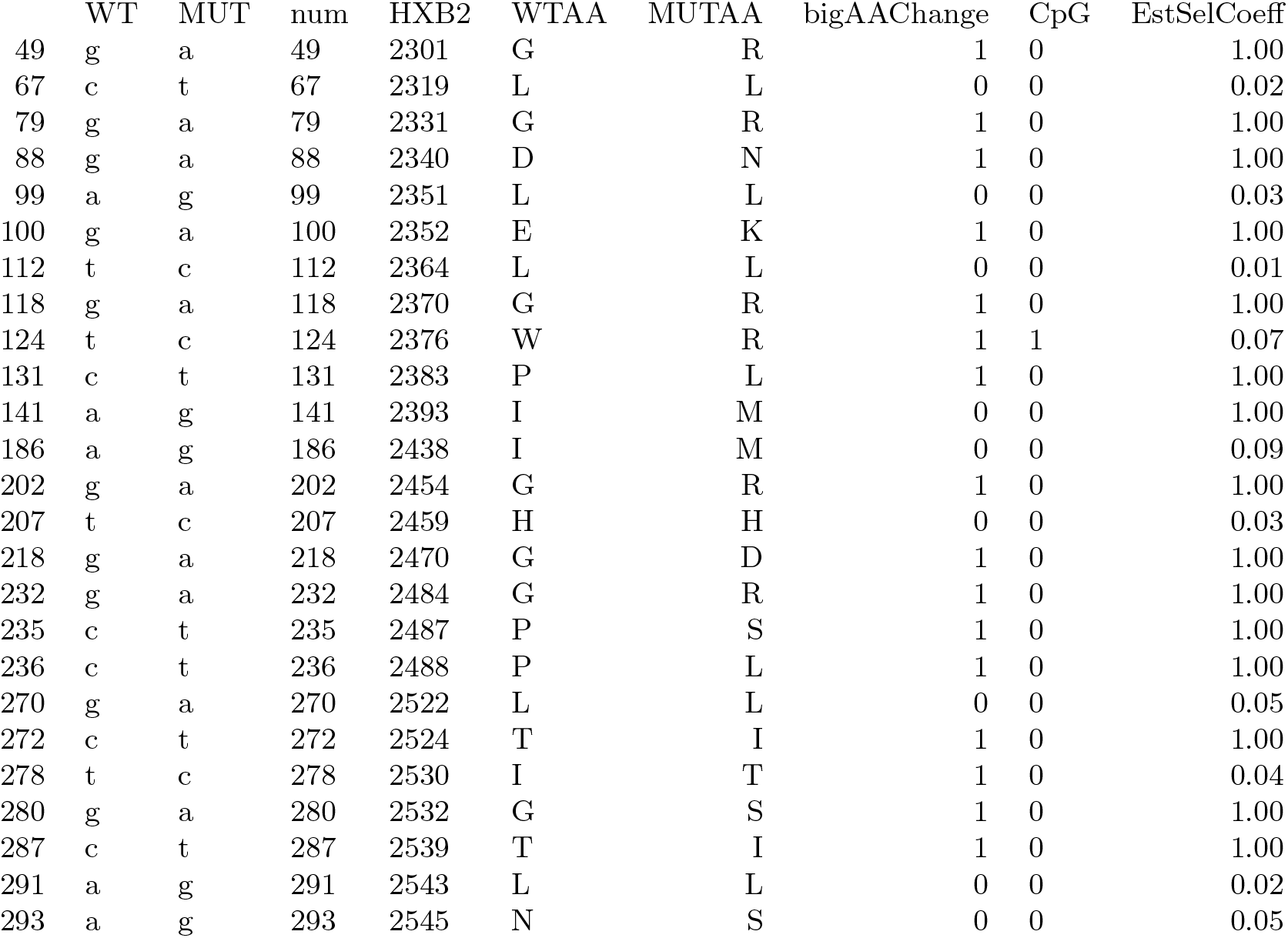
List of outlier sites with highest selection coefficients in *protease*. All sites were grouped in 9 groups, then the 5% highest selection coefficients were recorded in each group.

### 2.6 Relationship between mutation frequencies within patients and within the global subtype B epidemic

Next, we wanted to determine how well the observed within-patient mutation frequencies correspond with worldwide HIV mutation frequencies. All sequences in the Bacheler *et al* dataset belonged to HIV-1 subtype B, which is the most studied HIV-1 clade. We assembled a comparison set of HIV-1 subtype B sequences from treatment-naive patients using the Stanford HIV Drug Resistance database (HIVdb); this set contained 23,742 *protease* sequences and 22,785 reverse transcriptase sequences [47]. Fig 11 shows the correlation between average within-patient mutation frequencies from the 160 patients analyzed in this study and global mutation frequencies calculated from the HIVdb dataset. A high correlation coefficient was detected when comparing all 870 sites (Spearman’s rank correlation coefficient ρ = 0.68), showing concordance between mutation frequencies within patients and in the global subtype B epidemic. Similarly, Zanini *et al* [31] found that fitness costs were anti-correlated with subtype diversity (Spearman’s rank correlation coefficient ρ = -0.59)

## Discussion

Our fitness cost inference approach is based on the simple but highly powerful notion that mutation frequencies are in mutation-selection balance. We began by validating the approach. First, as expected, we found a clear separation of observed frequencies for synonymous, non-synonymous and nonsense mutations (Fig. 1). Second, we found that inferred costs of drastic amino-acid alterations were higher than those of non-drastic changes (Fig. 2). This matches biological knowledge, and has been observed when analyzing long-term evolution [48,49]. To the best of our knowledge this is the first report that physicochemical differences in amino-acids directly affect short term evolution as occurring during within-host evolution (but see [50]). These validations allowed us to focus on novel insights obtained by the method. First, we found that mutations that created new CpG dinucleotides were twice as costly as mutations that did not. Although it has been known for a while that CpG sites are depleted in genomes of HIV [42, 51, 52] and other RNA viruses [40–43], this is the first report suggesting strong selection operating against the de novo creation of CpG sites via mutation, to the extent that even one more CpG causes a fitness cost. Indeed, just recently it has been reported that HIV viruses with multiple CpGs in their genome are actively detected by an innate immune enzyme called ZAP, leading to inhibition of viral replication [44]. Our results hence suggest that this line of defence is particularly potent in driving the evolution of HIV, at the resolution of single nucleotide changes. Our next surprising finding was the substantial difference in fitness cost depending on which of the four nucleotides was altered. In particular, G→A and C→T mutations were two to seven times more costly than A→G mutations (discussed below). Thus, although we analyzed only a small part of the HIV genome using a dataset with limited sequencing depth, we succeeded in recovering and quantifying many known properties of mutational fitness costs, as well as discovering novel findings. Our data also allowed us to estimate parameters of DFEs, which will be useful for future studies on the evolutionary dynamics of HIV populations (Fig. 5, Table 2). Finally, we found that within-patient frequencies and global frequencies in the subtype B clade were very similar (Spearman’s rank correlation coefficient ρ = 0.68), suggesting that fitness costs are largely similar both within patients and across the pandemic.

### Comparison with other studies in viruses

In general, our results are consistent with those from a recent study on HIV-1 evolution by Zanini *et al* [31], based on a dataset described previously by the same authors [45]. Notably, both studies found a clear separation between synonymous and non-synonymous mutation frequencies, and these frequencies correlated well with global HIV diversity. Our study went on to find several novel insights. It should be noted that the proportion of lethal mutations estimated in our study (5.9%) is low compared to proportions from [31] and from *in vitro* studies on viral coding sequences (reviewed in [53]). For example, Sanjuan *et al* [22] found that 40% of random mutations in the RNA vesicular stomatitis virus were lethal. Similarly, a study by Rihn *et al* [54] of the HIV capsid found that 70% of non-synonymous mutations were lethal, which corresponds to around 47% of all mutations [54].

Several factors could explain why we found a lower proportion of lethal mutations as compared to other studies. First, even observed variants may represent inviable viruses, and for many sites in our dataset, the bootstrapped confidence intervals include lethality (data not shown). Second, we only considered transition mutations, whereas transversions may be more frequently lethal, as they are more often non-synonymous, more likely to lead to drastic amino acid changes, and more likely to create premature stop codons, due to the nature of the genetic code. Third, sequencing or amplification (PCR) errors may obscure our results. Many low-frequency variants in our dataset were only observed once, and it is possible that some of these were not true variants; we may thus have underestimated the percentage of lethal mutations. Fourth, we looked only at one gene, and this gene may have a different fitness landscape than other parts of the viral genome. Finally, different environments (*in vitro* vs *in vivo*) or different genetic backgrounds (usually one genetic background in the *in vitro* studies vs many in *in vivo* studies) may explain the observed differences. Future studies with more sequences and more sites will have better power to determine the true proportion of lethal mutations in HIV *in vivo*.

### High costs of G→A and C→T mutations

Among synonymous and non-synonymous mutations, we found G→A mutations to be two-and-a-half to seven times more costly than A→G mutations. C→T mutations were found to be two to five and a half times more costly than A→G mutations. We suggest three hypotheses to explain these initially surprising results: 1. They could be an artifact caused by spurious mutation rate estimates, 2. This could be due to mutation bias present in HIV genomes, 3. This could represent a form of APOBEC3 hypermutation. We’ll discuss these in the following paragraphs.

(1) *Mutation rates estimates*. We note that synonymous G→A mutations are present in our data at higher frequencies than A→G mutations (see Fig. 2. Naively, this would suggest that they are less costly. However, the G→A mutation rate is estimated to be so high that the observed frequencies are actually lower than expected, which, in our model translates to high costs. Synonymous C→T are equally frequent as A→G mutations, but, because they have a higher mutation rate, we conclude that they must be associated with higher costs. Hence, the mutation rate estimates are key to these results. We have two sources for mutation rate estimates from two very different studies, Abram *et al* [29, 38] (*in vitro estimates* from cell culture) and Zanini *et al* [31] (*in vivo* estimates based on accumulation of synonymous mutations). Notably, in both of these studies, G→A mutations occur at a higher rate than other transition mutations, although in the Zanini study this difference is less pronounced (Fig. 12). Using mutation rates from one or the other study does not change our findings qualitatively. If however G→A and C→T mutation rate estimates are overestimated in both studies, we cannot rule out that fitness costs of these mutations are lower than what we estimate.

(3) Second, the effect of costly G→A and C→T mutations might be related to a strong mutation bias in the HIV genome. G→A mutations are roughly five times more common than A→G mutations according to [29] and two and a half times more common according to [31] (Fig. 12 and Table 6). C→T mutations are twice as common as A→G mutations in according to both studies (see Table 6). Specifically, the G→A bias may have led, over long evolutionary timescales, to the well known A bias in the HIV genome [55,56]. Due to the strong mutation bias, sites at which having an A or G does not affect viral fitness would become A-biased over time. Thus, A sites would be enriched for (nearly) neutral sites, and G sites would be depleted of neutral sites, which could lead to G→A mutations being more costly, on average, than A→G mutations. A similar effect may be at play for C→T mutations, since here is also a T bias in the HIV genome, though it is not as strong as the A bias.

(3) *APOBEC3 hypermutation*. The effect of costly G→A mutations may be related to the activity of APOBEC3 enzymes, which hypermutate the HIV genome, leading to an increased proportion of G→A mutations [57–60]. We checked whether our sequences are dramatically affected by APOBEC3 hypermutation. Visual inspection of neighborjoining trees for each patient showed that there were no *pol* sequences that were hypermutated. This is probably because hypermutated viruses are mostly non-viable, and unlikely to show up when genetic material from viral particles is sequenced, which is what the current study is based on. However, APOBEC3 may also have a milder effect of slightly increasing the number of G→A mutations in the genome. The G→A mutations we observe could be linked to other G→A mutations in the genome, outside the sequenced region of *pol*. Together, these G→A mutations could be more deleterious than a single A→G mutation (which we use for comparison). This could explain why the observed G→A mutations in our study are more costly.

### Study limitations

One limitation of our study is that we focused on a small region of the HIV genome, namely 870 sites of the *pol* gene [61]. Because the patients in the Bacheler *et al* study were treated with a variety of antiviral treatments, we had to exclude drug resistance positions, as they would have been under positive selection in some of the patients. To study the costs of resistance mutations, it would be necessary to analyze samples from untreated patients [31]. Furthermore, is unknown how long the patients in our dataset were infected before samples were collected. If samples were taken soon after infection, genetic diversity in the viral population may have been low, and frequencies of some mutations may have been lower than the expected f = *u/s*, resulting in overestimates of the selection coefficients. A second limitation is that we assumed one mutation rate for all A→G mutations, and one rate for all C→T mutations, etc. However, evidence exists that mutation rates vary along the genome, which would mean that selection coefficient estimates for individual mutations may be unreliable [62,63].

Finally, our *in vivo* frequency-based approach did not allow us to study epistatic interactions between mutations. Recent work on HIV, however, shows that epistatic interactions may be important. For example, such interactions play a role in determining the mutational pathway that the virus uses to escape cellular immunity [64] and to develop drug resistance [25, 65, 66]. It is currently unclear how the costs of mutations as determined in this study depend on their genetic background and further studies need to be designed that combine the strengths of our approach to study costs of virtually all mutations *in vivo*, with the strengths of other approaches to study epistatic effects between common mutations.

### Outlook

The current study should be seen as a proof of concept of our *in vivo* frequency-based approach. Our results demonstrate the power of analyzing mutant frequencies from *in vivo* viral populations to study the fitness effects of mutations. We hope that soon this method will be applied to the entire HIV genome and the genomes of other fast-evolving microbes. For HIV specifically, we expect that patient samples with high viral loads will be sequenced much more deeply than in any of the studies analyzed in this article. Transversion mutations can then be analyzed in addition to transition mutations. Such a dataset will allow us to get a more fine-grained and precise picture of the costs of mutations at individual sites across the entire HIV genome, including for mutations in other genes and non-coding regions of the virus and for drug resistance mutations in *pol* and elsewhere. Because our method makes it possible to estimate *in vivo* costs, the results will contribute to our understanding of drug resistance evolution and immune escape and may also contribute to vaccine design.

## Methods

### Description of the data/filtering

We used sequences from a dataset collected by Bacheler *et al*. [61], a study that focused on patients in three clinical trials of different treatments, all based on efavirenz (a non-nucleoside *RT* inhibitor) in combination with NRTIs (nucleoside *RT* inhibitors) and/or *protease* inhibitors. The treatments in this study were not very effective, in part because some patients were initially prescribed monotherapy, which almost always lead to drug resistance, and in part because patients had previously been treated with some of the drugs, so their viruses were already resistant to some components of the treatment. Viral loads in these patients were typically not suppressed, which made it possible to sequence samples even during therapy. We have previously used part of this dataset to study soft and hard selective sweeps [35].

The Bacheler *et al*. [61] samples were cloned and Sanger-sequenced. For each patient, all available sequences were treated as one sample, even when they came from different time points. Patients with less than five sequences were excluded from the analysis, leaving us with a median of 19 sequences per patient for 160 patients (3,572 sequences in total). Sequences were 984 nucleotides long and were composed of the 297 nucleotides that encode the HIV *protease* protein and the 687 that encode the beginning of RT. We excluded 75 drug resistance–related sites [67] and 39 *protease* sites that overlap with gag, leaving 287 synonymous, 555 non-synonymous and 28 nonsense mutations, for a total of 870 sites. Sequences were retrieved from Genbank under accession numbers AY000001 to AY003708.

### Calculation of mutation frequencies

To identify mutations, we compared the sequences to the HIV-1 subtype B reference sequence, also known as the HXB2 sequence (Genbank accession K03455). We will refer to this reference sequence as the wildtype (WT) or ancestral sequence. To make sure that mutations in founding viruses with which patients got infected not skew our results, we added a filtering step. For each patient, sites are only included if all sequences from the first sampling time point for that patient carry the same nucleotide as the reference B WT sequence. This filtering step removed 6% of the data. We only considered transition mutations (A↔G and C↔T), excluding transversion mutations. For example, for a site with an A in the reference sequence, the frequency of a transition mutation was calculated for each patient as the number of sequences with a G divided by the number of sequences with a G or an A. Sequences with a C or a T were thus not considered at all if the reference sequence had an A in that position. In addition, if, in a given sequence, there was more than one mutation in a triplet, this triplet was removed for that specific sequence, so that all mutations could be clearly classified synonymous, non-synonymous or nonsense. Occasionally this meant that a sample from a patient had to be excluded for a given site, so for some mutations we had fewer than 160 frequencies to analyze.

Selection coefficients were estimated for each mutation by dividing the nucleotide-specific mutation rate by the observed average frequency (based on the mutation-selection balance formula f = *u/s*). We used mutation rates as estimated by Abram *et al*. [29, 38].

### Sliding window approach to determine location effect

This analysis aims to determine whether sites that are in close proximity to each other have more similar fitness costs than expected. If the window size is 10, then we first consider the first 10 non-synonymous sites in the *pol* gene and we calculate the mean fitness effect of the mutations in that window (window mean). We then slide with step size 1 to sites 2 to 11 and again calculate the window mean fitness effect etc. In this manner we slide from the beginning to the end of the sequence and once we have all window means, we calculate the variance of the window means. If high cost sites are clustered spatially, than the mean fitness is high in some windows but low in others and the variance of the window means will be relatively high. We compared the variance of window means with the null expectation of no spatial clustering. To obtain a null expectation, we randomized the location of all positions, while keeping the sequence the same (e.g., each non-synonymous G-A mutation would be swapped with another non-synonymous G-A mutation). For the resulting randomized datasets we also calculated the variance of the window means. We then compared the range of variances obtained from 1000 randomizations with the variance from the real data. For synonymous sites, the observed variance of window means was never significantly higher than the variance of window means of randomized datasets, for a wide range of window sizes (2-100), which shows that there is no evidence for any location effect for synonymous sites, in other words, there are no stretches of low or high fitness cost mutations.

For non-synonymous sites, we found that the variance of window means for the real data was often higher than the variance of window means for the randomized data, which suggests that, for non-synonymous sites, there are stretches of the *pol* gene with higher fitness costs and stretches with lower fitness costs. We hypothesized that this was due to the fact that two neighboring nucleotides within a codon, will affect the same amino acid, and if that amino acid is important for the fitness of the virus, then mutations at both of the nucleotides will be particularly costly. To test this, we did a randomization test where we kept codons in tact, but randomized their location. For example, a codon that encodes for asparagine could be swapped with another codon that encodes for asparagine. We found that after this codon by codon randomization, we find the same variance of window means as we find in the original dataset. This shows that the location effect we see is mostly due to neighboring sites within codons.

### Generalized linear model analysis

Using a *generalized linear model* (GLM), we predicted mutant frequencies for certain categories of mutations (e.g., synonymous, non-CpG-forming, A→G mutations) and then used the mutation-selection formula (f = *u/s)* to predict the costs of these groups of mutations (see Fig. 10). Specifically, we fit a GLM where the response variable is whether a given nucleotide is WT or mutant, and this response variable is assumed to follow a binomial distribution, using the *glm* package in the *R* language [68]. The model we fit includes the nucleotide in the consensus sequence, its experimentally determined SHAPE value [39], whether or not the position was in the *RT* protein and the types of changes resulting from a transition at that position. These changes included whether a transition was non-synonymous, lead to a drastic amino acid change or formed a new CpG site. We used the following groups of amino acids and assumed that a change from one group to another was ‘drastic’: positive-charged (arginine (R), histidine (H), lysine (K)), negative-charged (aspartic acid (D) and glutamic acid (E)), uncharged (serine (S), threonine (T), asparagine (N) and glutamine (Q)), hydrophobic groups (alanine (A), isoleucine (I), leucine (L), phenylalanine (F), methionine (M), tryptophan (W), tyrosine (Y) and valine (V)), the special amino acids (cysteine (C), glycine (G) and proline (P)). We also fit interactions between the ancestral nucleotides, whether a transition was non-synonymous, and whether the transition formed a CpG site.

Note that for the GLM, actual counts were considered as opposed to frequencies. That is, if we have 20 sequences for patient 1, and at a given nucleotide, we observe 2 As and 18 Gs, we used those counts. This approach automatically gives more weight to patients for whom we have more sequences. Each position in each sequence from each patient was treated as an independent observation.

The GLM coefficients reported in table 1 can be used to predict the probability that a mutation is observed at a given site. For example, the intercept = (—5.2) means that a synonymous, non-CpG-forming mutation in *protease* at a site with A as WT has an probability of *exp*(−5.2) = 0.055 to be mutated, so its predicted frequency is 0.055. For a similar site that has T as WT, we need to add 0.013 to the exponent and find a probability of *exp*(−5.2 + 0.013) = 0.056.

To explicitly test whether two categories of mutations with different mutation rates had different selection coefficients, we used a one-sided two-sample Wilcoxon test (also known as a Mann-Whitney test). This was necessary because a GLM can only test whether a mutant of a certain category is more likely to be present than a mutant of another category (i.e., has a higher frequency). We were interested, however, in whether a mutant of a certain category is more costly than a mutant of another category. For example, synonymous C→T mutations occur at a similar frequency as synonymous, non-CpG forming A→G mutations (see Table 1, line 5), but because their mutation rates are quite different, we estimate that their costs are different. (see Fig. 10).

### Outlier analysis

We grouped the sites first in nine groups according to the GLM results and then listed outliers (5% highest selection coefficient values) in each group.

The groups used were:

- synonymous, non-G, non-CpG
- synonymous, non-G, CpG
- synonymous, G
- non-syn, A or T, no-CpG, no-drastic AA change
- non-syn, A or T, CpG, no-drastic AA change
- non-syn, A or T, no-CpG, drastic AA change
- non-syn, A or T, CpG, drastic AA change
- non-syn, C or G, no-CpG, no-drastic AA change
- non-syn, C or G, no-CpG, drastic AA change

### Estimating a gamma distribution to fit the distribution of fitness effects

We fit a gamma distribution to the DFE (based directly on averaged frequencies at 870 sites and the mutation-selection balance formula *f* = *u/s)*. Transitions that were never observed (frequency of 0) were not considered when fitting the gamma distribution. The most likely shape and scale parameters for the data were found using the subplex algorithm implemented in the R package nloptr [69] (see Table 2). Bootstrapped confidence intervals were created by resampling the data with replacement and re-estimating the gamma distribution parameters. Selection coefficients were estimated using the mutations rates given in Abram *et al*. [29,38] and Zanini *et al*. [31].

### Comparison with the global epidemic

A large HIV-1 sequence dataset was retrieved from the HIVdb (http://hivdb.stanford.edu/pages/geno-rx-datasets.html) [47]. This dataset contains a single sequence per patient. *Protease* and *RT* sequences were downloaded in separate files. Sequences that met the following criteria were included in the analysis: treatment-naive host status and classification as HIV-1 subtype B. In total, 23,742 *protease* and 22,785 *RT* sequences were collected. Average mutation frequencies for each site were calculated as explained above (e.g., including only transitions, excluding triplets with more than one mutation). Spearman’s rank correlation coefficient *(ρ)* was used to quantify the correlation between within-patient and global mutation frequencies.

### Additional datasets

In order to test how transferable our method is, we repeated parts of our analysis with the Zanini *et al*. dataset [45] and the Lehman *et al*. dataset [46].

The Zanini [45] samples came from nine patients. There were multiple samples per patient (72 samples in total), typically collected at least a few months apart. Thus we followed Zanini *et al* in treating those samples as if they were completely independent. The sequencing method used was Illumina. We downloaded mutation frequencies for each sample (http://hiv.tuebingen.mpg.de/data/) and averaged frequencies across all 72 samples. The Zanini data cover the whole HIV genome, but we only considered the regions that overlap with the Bacheler data [61]. In addition, the Zanini data [45] contain sequences for different HIV subtypes (B, C and CRF01-AE); we only considered sites that were conserved between subtypes B, C and CRF01-AE and excluded resistance related sites so that 758 sites were analyzed. Mean mutation frequencies for all sites, ordered by mutation frequency are shown in Figure S1. The distribution of fitness effects is shown in Figure S3 and the estimated gamma distribution parameters in Table 5.

**Table 4:**
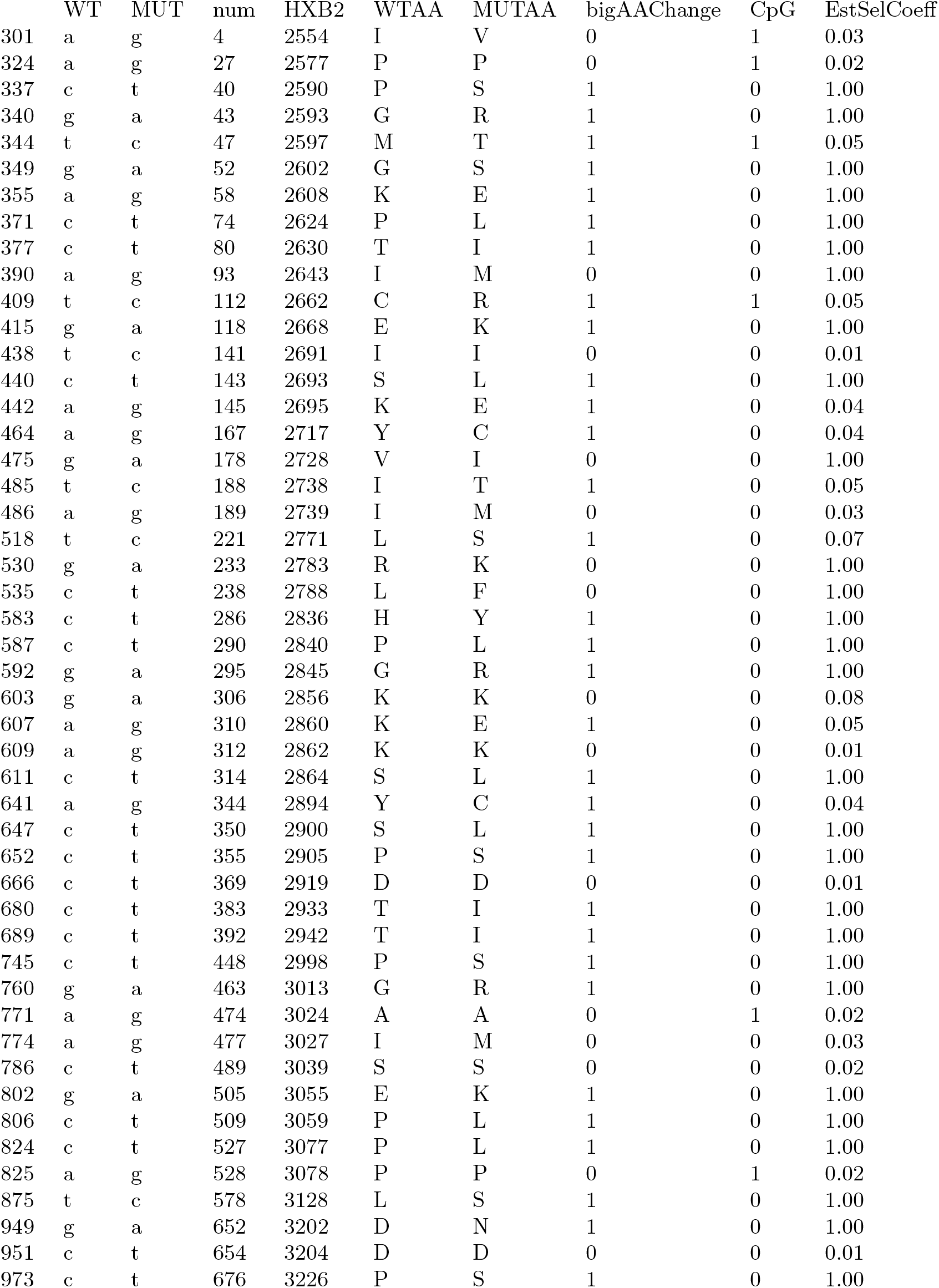
List of outlier sites with highest selection coefficients in reverse transcriptase. All sites were grouped in 9 groups, then the 5% highest selection coefficients were recorded in each group.

The Lehman samples were 454-sequenced. The samples were collected at seroconversion and one month later, but we only included the time point one month after seroconversion in our analysis, as we expected that the samples from the earliest time point would contain almost no genetic diversity. The sequences span approximately 540 sites in the *RT* protein. The Lehman data [46] contained HIV subtypes B, C and A; we only considered sites that were conserved between subtypes B, C and A and excluded resistance related sites, so that 415 sites were analyzed. Mean mutation frequencies for all sites, ordered by mutation frequency are shown in Figure S2. The distribution of fitness effects is shown in Figure S4 and the estimated gamma distribution parameters in Table 5. The Lehman dataset [46] was downloaded from the NCBI website using accession number SRP049715 (www.ncbi.nlm.nih.gov/sra/?term=SRP049715).

## Acknowledgments

The authors wish to thank Dmitri Petrov, Arbel Harpak, David Enard, Nandita Garud, Alan Bergland and Ryan Taylor for helpful discussions; Richard Neher, Fabio Zanini, Adam Eyre-Walker and an anonymous reviewer for comments on earlier versions of the manuscript; Scott Roy for help aligning the Lehman sequences.

## Supporting information

S1 File. Mutation frequencies and estimated selection coefficients from the Bacheler [61] dataset.

**Fig S1:**
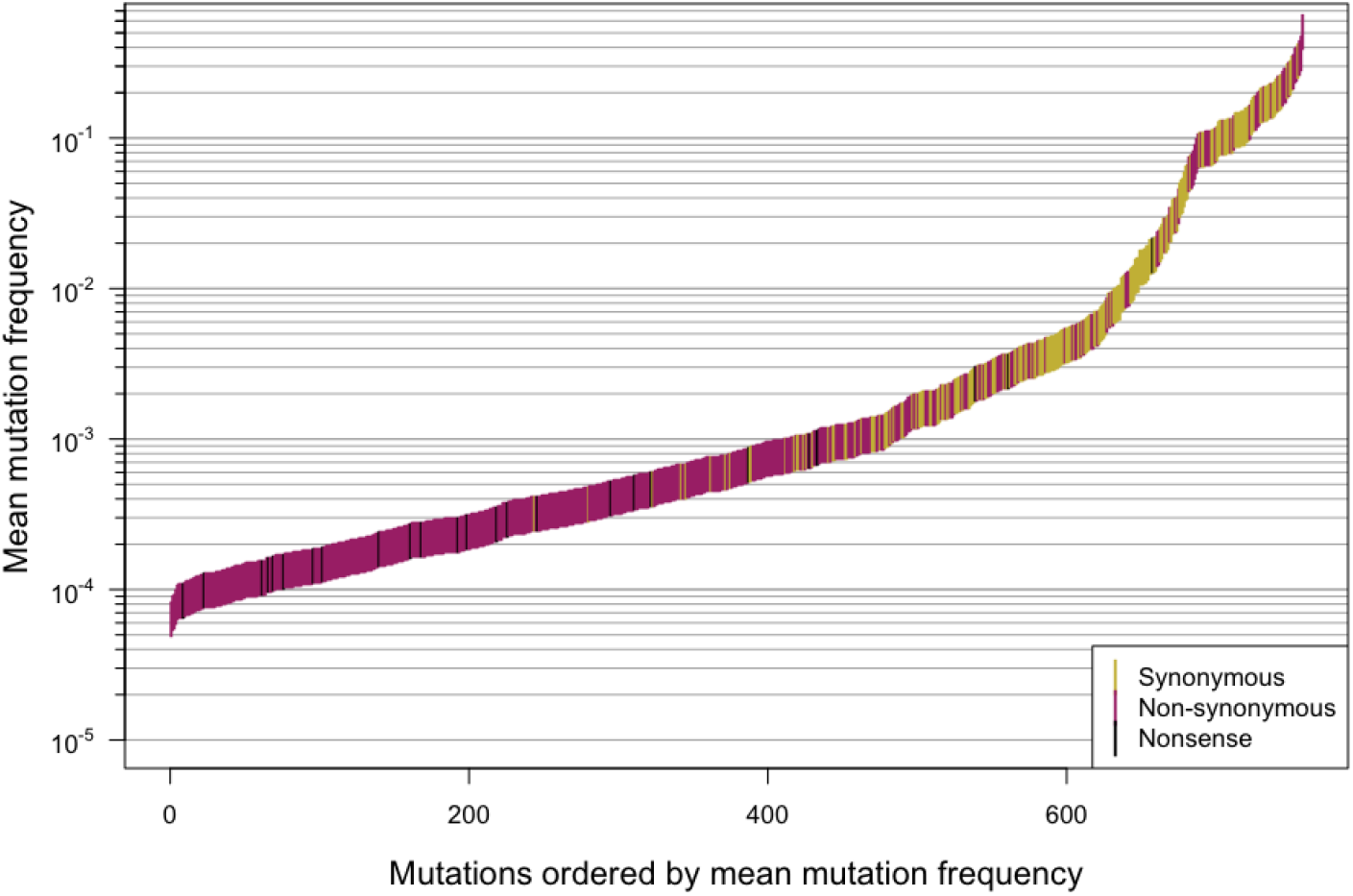
Mutation frequency for the Zanini dataset [45]. Mutation frequency for 758 *pol* sites from the Zanini dataset [45], ordered by mutation frequency.

**Table S1:**
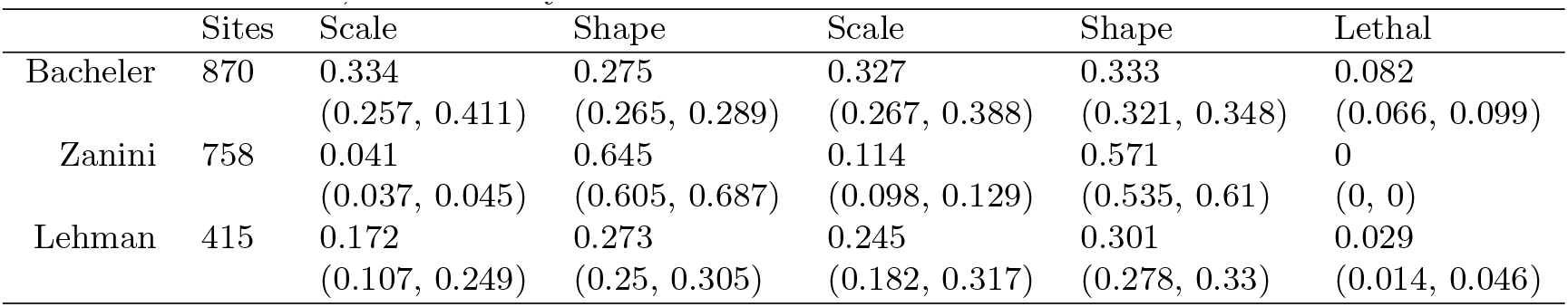
Parameters for the gamma distribution of fitness costs for the Bacheler, Zanini and Lehman datasets [45,46,61]. Parameters for the gamma distribution of fitness costs for *pol* mutations based on mutation frequencies the Bacheler, Zanini and Lehman datasets, reflecting scale (κ) and shape (θ). The “fraction lethal” is the fraction of the mutations that had a mean frequency smaller than or equal to the mutation rate, so that they are estimated to be lethal.

**Fig S2:**
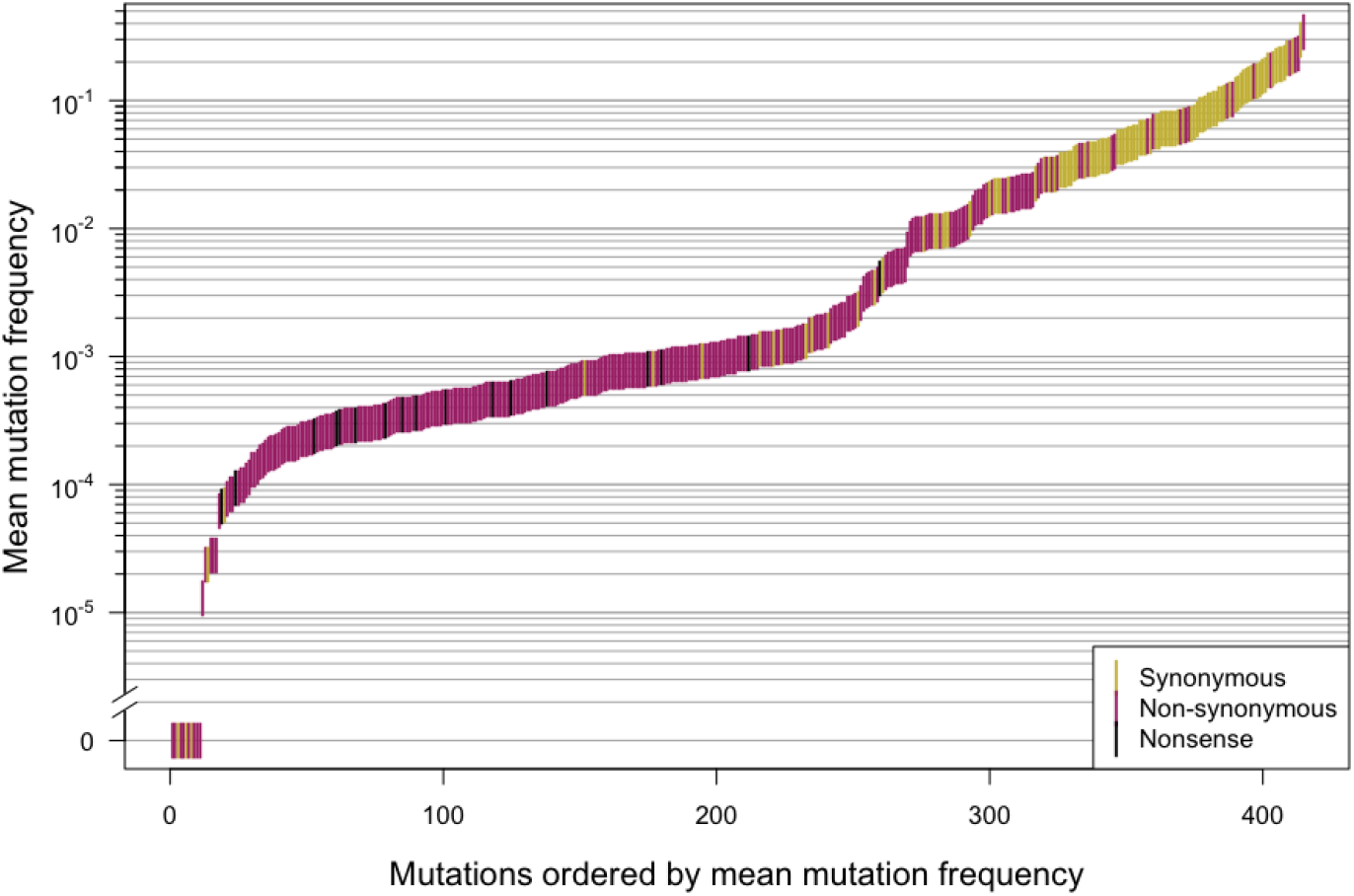
Mutation frequency for the Lehman dataset [46]. Mutation frequency for 621 reverse transcriptase sites from the Lehman dataset [46], ordered by mutation frequency.

**Fig S3:**
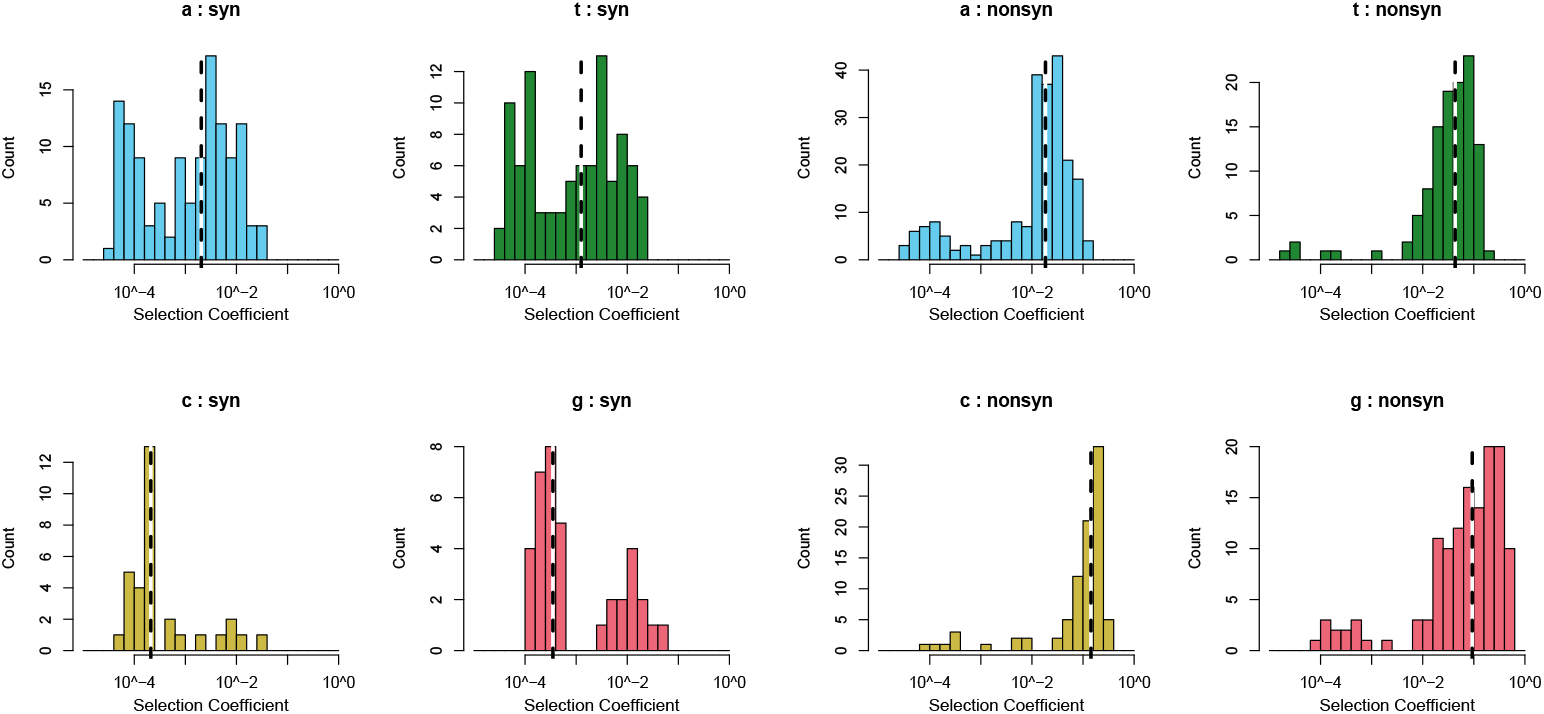
Distribution of fitness costs for the Zanini dataset [45]. Distribution of fitness costs for non-synonymous and synonymous mutations for the Zanini dataset [45]. Nonsense mutations are included in the non-synonymous mutations. Note that the scale of the y-axis differs between the graphs.

**Fig S4:**
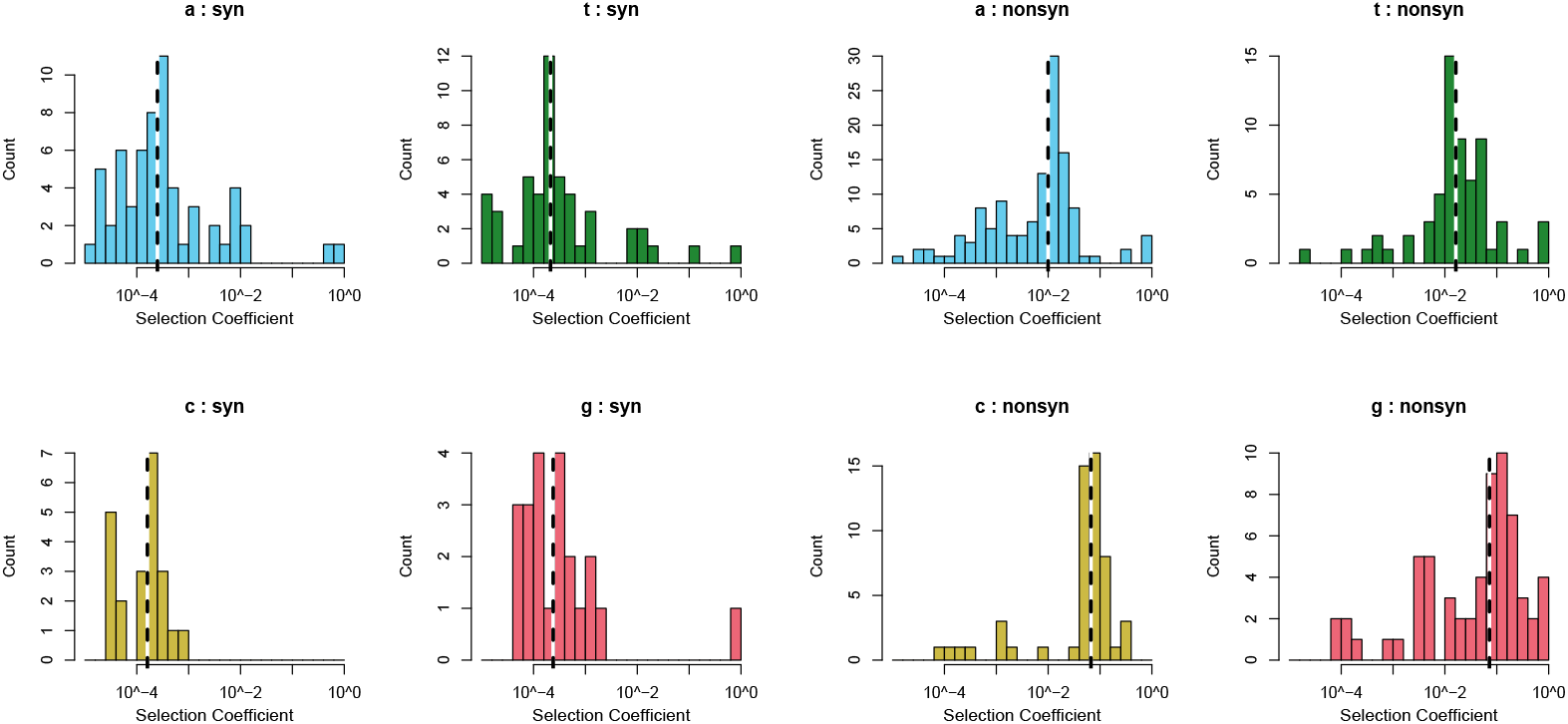
Distribution of fitness costs for the Lehman dataset [46]. Distribution of fitness costs for non-synonymous and synonymous reverse transcriptase mutations from the Lehman dataset [46]. Nonsense mutations are included in the non-synonymous mutation category. Note that the scale of the y-axis differs between the graphs.

**Fig S5:**
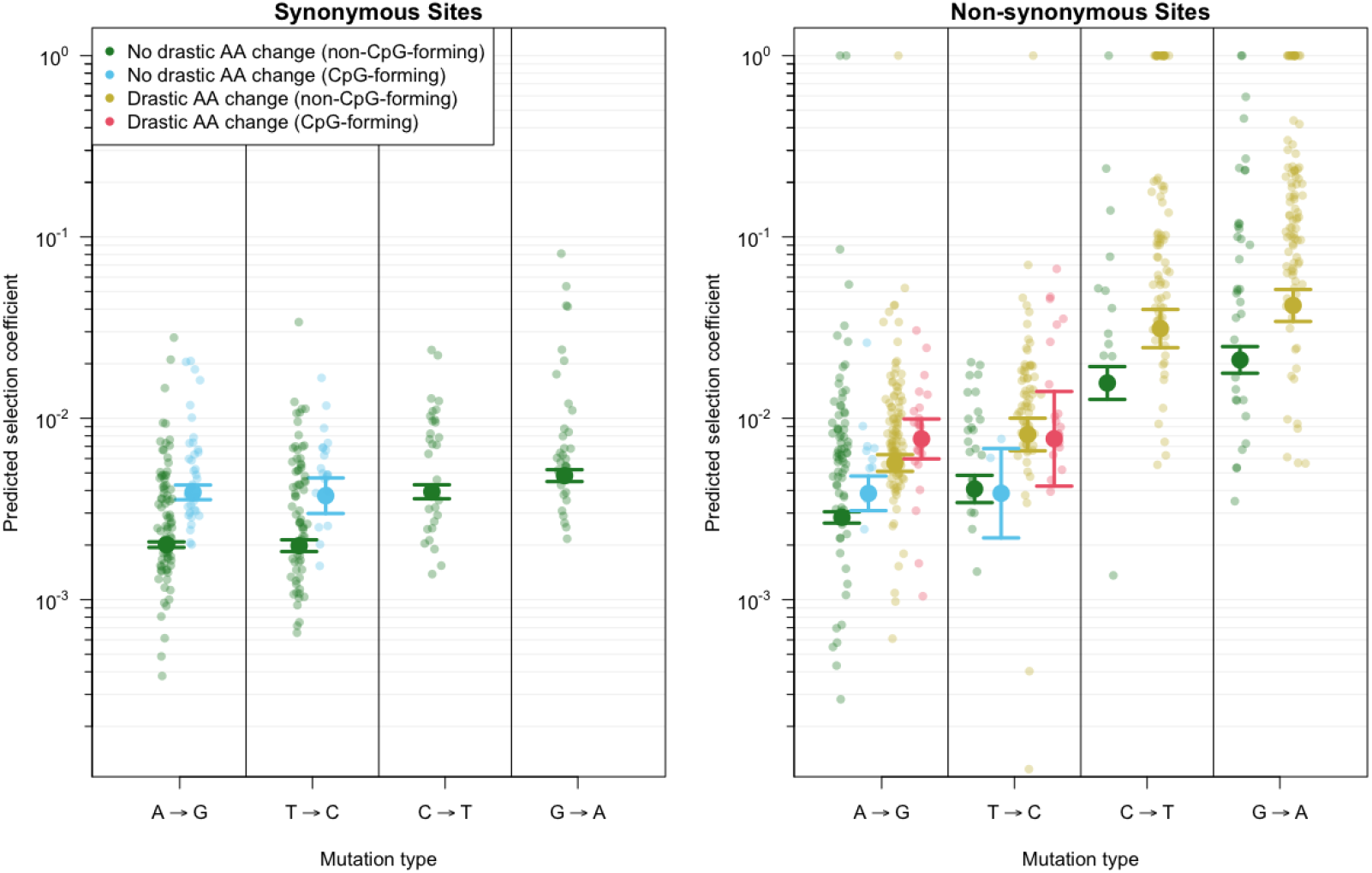
Estimated selection coefficients for different mutation classes. Selection coefficients for transitions at every nucleotide site in the *pol* sequence show that CpG-forming mutations are more costly than non-CpG-forming mutations and that mutations that involve a drastic amino acid change are more costly than mutations that do not. Selection coefficients were estimated using a generalized linear model and sequence data from 160 HIV-infected patients. Shown are predicted selection coefficients for synonymous (left) and non-synonymous (right) mutations that do not involve a drastic amino acid change and either create CpG sites (blue) or do not (green). For non-synonymous mutations, predictions are also shown for mutations that do involve drastic amino acid changes and either create CpG sites (light red) or do not (yellow).

**Fig S6:**
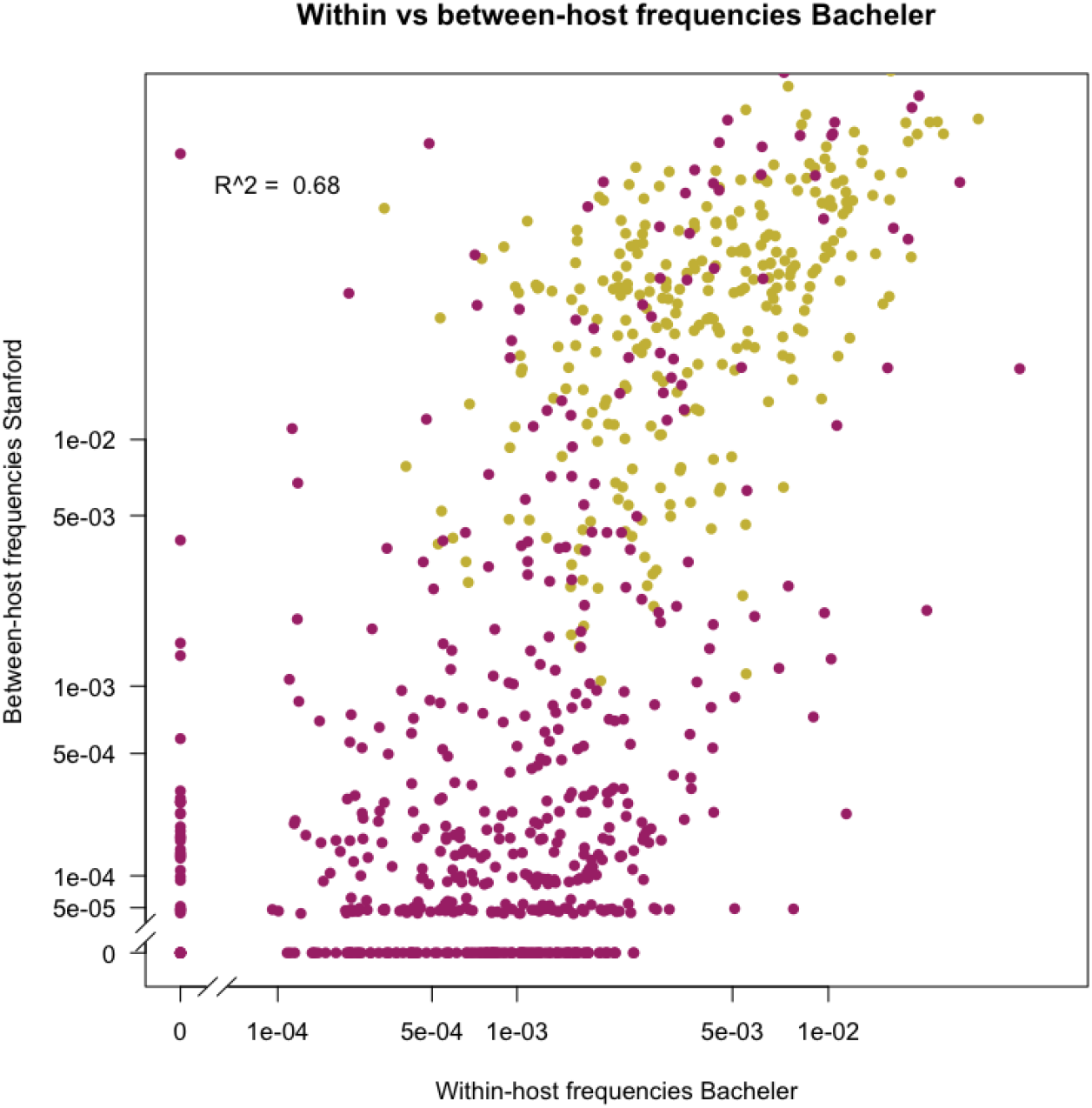
Correlation of within-patient mutation frequencies and global between-patient subtype B mutation frequencies. A correlation (Spearman’s rank correlation coefficient ρ = 0.68) exists between average *pol* mutation frequencies at the within-patient level (in the 160 patients analyzed in this study) and mutant frequencies in the global subtype B epidemic (23,742 *protease* and 22,785 reverse transcriptase consensus sequences from the HIVdb [47]). Values shown on a log scale. Non-synonymous mutations are shown in dark pink, synonymous mutations in yellow.

**Fig S7:**
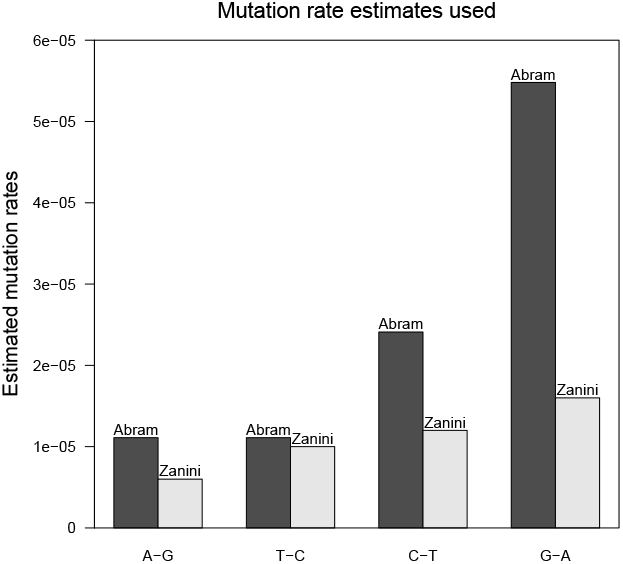
Mutation rate estimates per replication from Abram *et al*. [62] as calculated by Rosenbloom *et al*. [38] and mutation rate per day from Zanini *et al*. [31].

**Table S2:**
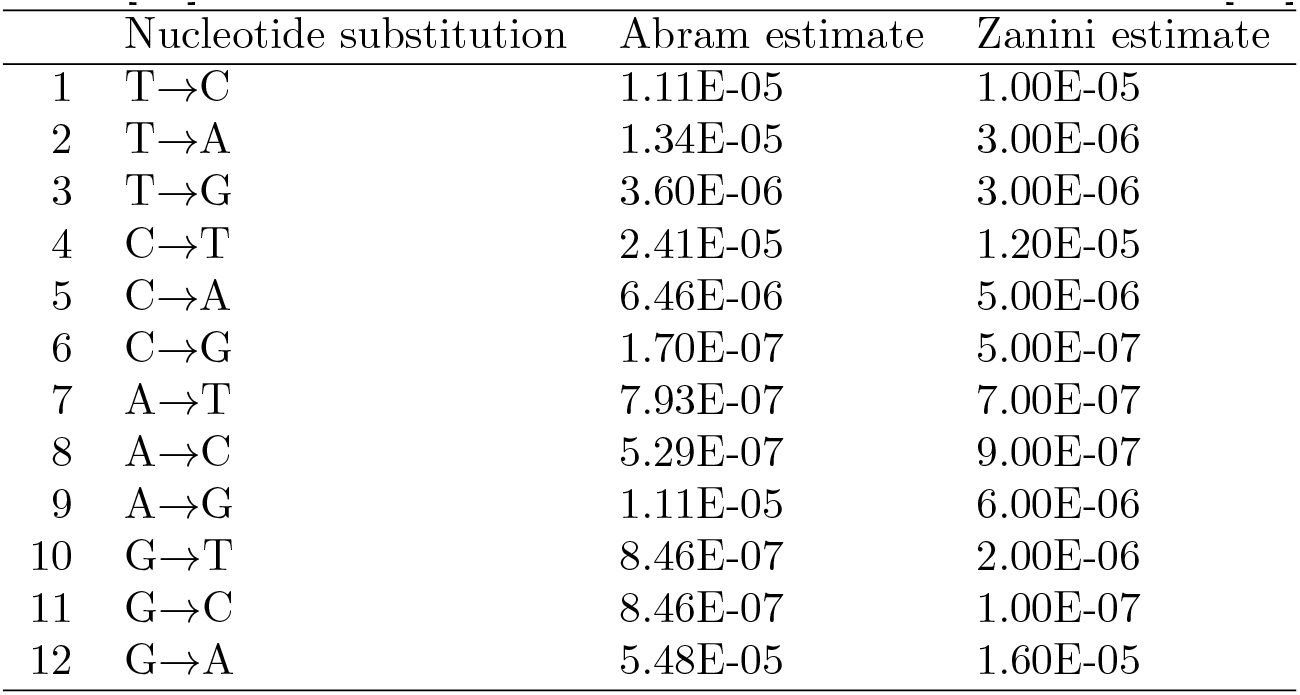
Mutation rate estimates per replication from Abram *et al*. [62] as calculated by Rosenbloom *et al*. [38] and mutation rate per day from Zanini *et al*. [31].

